# DMRT1 regulation of *TOX3* modulates expansion of the gonadal steroidogenic cell lineage

**DOI:** 10.1101/2022.07.29.502037

**Authors:** Martin A. Estermann, Andrew T. Major, Craig A. Smith

## Abstract

Vertebrate gonads comprise three primary cell types, germ cells, steroidogenic cells, and supporting cells. The latter are the first cell type to differentiate in the embryonic gonad and direct the formation of other somatic lineages. During gonadal sex determination, the supporting cell lineage differentiates into Sertoli cells in males and pre-granulosa cells in females. In the chicken embryo, the molecular trigger for Sertoli cell differentiation is the Z-linked gene DMRT1. Recently, single cell RNA-seq data indicate that that chicken steroidogenic cells, derive from differentiated supporting cells. This differentiation process is achieved by a sequential upregulation of steroidogenic genes and down-regulation of supporting cell markers. The exact mechanism regulating this differentiation process remains unknown. We identified the gene *TOX3* as a novel transcription factor expressed in embryonic Sertoli cells of the chicken testis. *TOX3* knockdown in males resulted in increased *CYP17A1* positive Leydig cells. *TOX3* over-expression in male and female gonads resulted in a significant decline in *CYP17A1* positive steroidogenic cells. *TOX3* expression is negatively regulated by estrogens *in vivo*, but not induced during masculinization induced by estrogen inhibition. *In ovo* knock-down of the testis determinant, *DMRT1*, in male gonads resulted in a down-regulation of TOX3 expression. Conversely, DMRT1 over-expression caused an increase in *TOX3* expression. Taken together, this data indicates that DMRT1 regulation of *TOX3* modulates expansion of the steroidogenic lineage, either directly, via cell lineage allocation, or indirectly via signaling from the supporting to steroidogenic cell populations.

## Introduction

During early embryonic development, the gonads typically differentiate into either testes or ovaries. In mammals, the testicular developmental pathway is initiated by the expression of the Y-linked gene *Sry* in the supporting cells, triggering the upregulation of thousands of genes over a short period of time and resulting in the differentiation of the supporting cells into Sertoli cells (1-3). One of the transcriptional targets of Sry is *Sox9*, an SRY-related HMG box family member, which is crucial in Sertoli differentiation (1, 4-7). Sertoli cells then direct differentiation of other testicular cell types (8). In the absence of this masculinizing signal, gonads differentiate into ovaries through the stabilization of the Rspo1/Wnt/β-catenin signalling pathway (9-11).

In the chicken model, the molecular trigger for Sertoli cell differentiation is the Z-chromosome linked gene, *DMRT1* (12). *DMRT1* is un-related to *Sry*, and it encodes a zinc-finger like transcription factor (13-16). Despite the differing gonadal sex determination triggers in mammals and birds, the genetic regulation of gonadal development and sexual differentiation is largely conserved. This includes the supporting cell markers *DMRT1, AMH* and *SOX9*, in the avian testis and the markers *WNT4/RSPO1, aromatase* and *FOXL2*, in the ovary (17-21). Recently, chicken single-cell RNA-seq data indicates that, although gonadal cell types are conserved, their developmental origin is not (22, 23). In the mouse embryo, the supporting cell lineage derives from the coelomic epithelium (24-26). In chicken, the coelomic epithelium gives rise to gonadal epithelium and interstitial cells (23). The supporting cells derive from a *DMRT1, PAX2, OSR1* and *WNT4* positive pre-existing mesenchymal population (23). Additionally, the scRNA-seq data strongly suggest that the steroidogenic cells derive from differentiating supporting cells (Sertoli and pre-granulosa cells) (23). This differentiation process is achieved by a sequential upregulation of steroidogenic genes, followed by the downregulation of supporting cell markers (23). Due to the novelty of this discovery in the chicken, the exact mechanism regulating this differentiation process remains unknown.

To expand our understanding of normal and aberrant gonadal development and differentiation, it is essential to identify novel regulators of ovarian and testicular development. Previously our laboratory identified *TOX3* (*TOX High Mobility Group Box Family Member 3*) as a novel gene expressed in chicken Sertoli cells (23). In mouse, *Tox3* is a high mobility group (HMG) box transcription factor predominantly expressed in the brain, where it plays a protective role inducing an anti-apoptotic response, interacting with CBP/CREB or CBP/CITED1 (27, 28). TOX3 acts as a transcriptional activator upregulating (directly or indirectly) a large number of genes involved cell proliferation, migration, mammary gland development and breast cancer (29, 30). Additionally, DNA variants in the *TOX3* locus have been associated with polycystic ovarian syndrome (PCOS) in several human populations, a syndrome associated with higher levels of androgens, (31-34). However, the exact functional mechanism of *TOX3* in this disease or in the gonadal context is unclear. In this manuscript we characterized the expression pattern of *TOX3* in the developing chicken gonad, focusing on how it is regulated and its potential role in gonadal supporting and steroidogenic cell differentiation. Our data support a model in which TOX3 modulates the steroidogenic cell population differentiation, and its dysregulation may underlie increased steroidogenic capacity, leading to PCOS.

## Results

### TOX3 is expressed in a subset of Sertoli cells after the onset of gonadal sex differentiation

Previous gonadal bulk RNA-seq performed in our laboratory showed that *TOX3* expression is sexually dimorphic at the onset of gonadal sex differentiation (Embryonic day 6, E6 / HH stage 29), being upregulated in male gonads (Fig. 1A) (35, 36). To validate these results, *TOX3* gonadal expression was quantified by qRT-PCR before E4.5 (stage 24), at the onset E6.5 (stage 30) and after E8.5 (stage 34) the onset of gonadal sex differentiation. *TOX3* was lowly expressed in both male and female gonads at E4.5 (Fig. 1B). *TOX3* mRNA expression was upregulated in developing male gonads at E6.5/stage 30 and E8.5/stage 34, whereas it remained low in female gonads (Fig. 1B). This data is consistent with the RNA-seq results. To define the spatial expression pattern of *TOX3* in embryonic gonads, whole mount *in situ* hybridization (WISH) was performed on male and female urogenital systems at E4.5 (stage 24), E6.5 (stage 30) and E8.5 (stage 34). Results showed positive staining in male gonads, but not in females, after the onset of gonadal sex determination (E6.5), consisting with qRT-PCR data (Fig. 1C). WISH gonadal transverse sections showed that, in males, *TOX3* was expressed in the developing seminiferous cords of the gonadal medulla (Fig. 1D).

**Fig. 1.**
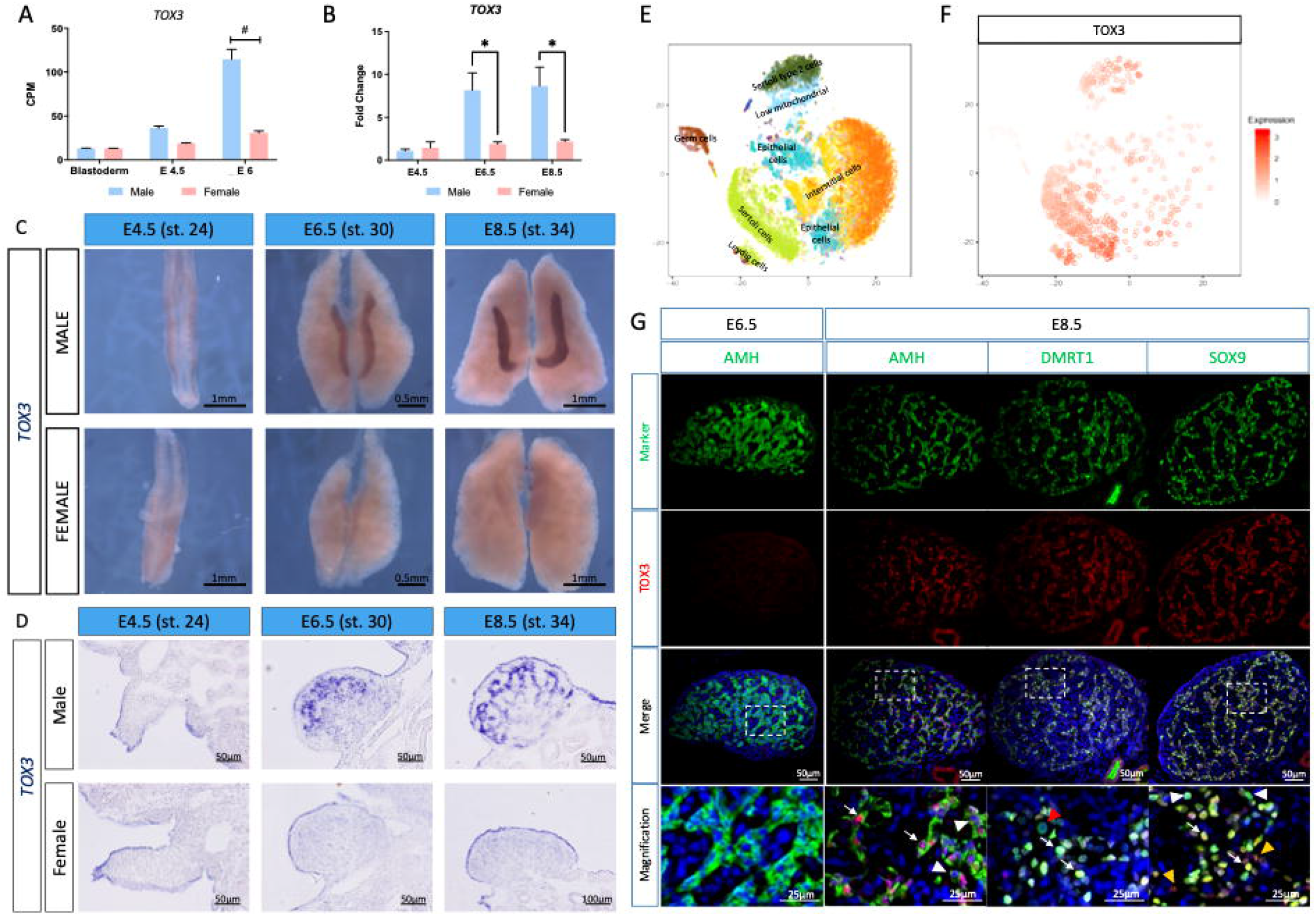
TOX3 is upregulated in chicken supporting cells after the onset of sex differentiation. (A) *TOX3* mRNA expression levels from bulk gonadal RNA-seq in count per million (CPM) at blastoderm stage, E4.5 and E6. # = false discovery rate (FDR) <0.001. (B) *TOX3* mRNA expression by qRT-PCR, relative to *β-actin* and normalized to E4.5 male. (Bars represent Mean ± SEM, n=6. * = adjusted p value <0.05. Multiple t-test and Holm-Sidak post-test. (C) *TOX3* mRNA expression by whole mount *in situ* hybridization in the urogenital system. (D) Sections of the *TOX3* whole mounts *in situ* hybridizations. (E) t-SNE plot of all gonadal male cells sequenced, color-coded by cell type. (F) Normalized expression of TOX3 on a t-SNE visualization of all male gonadal chicken cells. (G) E6.5 and E8.5 testicular immunofluorescences for Sertoli cell markers (AMH, DMRT1 and SOX9) and TOX3. White arrows indicate colocalization of TOX3 with the Sertoli cell markers. Dashed box indicates the magnified area. White arrowheads indicate AMH, SOX9 or DMRT1 positive, TOX3 negative cells. Yellow arrowheads indicate TOX3 positive SOX9 negative cells. Red arrowheads indicate DMRT1 positive germ cells.

To identify the specific testicular cell types expressing *TOX3*, our previous chicken testis single-cell RNA-seq data was scrutinized (23). A t-SNE containing E4.5, E6.5, E8.5 and E10.5 whole testis samples was generated, identifying the main testicular cell populations (Fig. 1E). *TOX3* expression was mainly restricted to the Sertoli cell lineage, as well as in the Sertoli-Leydig cell cluster (Fig. 1F). To confirm this expression pattern, immunofluorescence was performed in E6.5 and E8.5 testicular sections. Although *TOX3* mRNA expression was detected at E6-E6.5, TOX3 protein was not detected at this stage, indicating a delay in TOX3 translation or possibly expression below the level of detection with the antibody (Fig. 1G). At E8.5, TOX3 protein showed nuclear localization, as expected of a transcription factor, and was detected inside the testicular cords, consistent with the mRNA expression. TOX3 was expressed in supporting cells, colocalizing with SOX9, AMH and DMRT1 (Fig. 1G, white arrows), but not in the DMRT1^+^ germ cells (Fig. 1G, red arrowhead). Some AMH^+^, SOX9^+^ supporting cells were negative for TOX3 (Fig. 1G, white arrowheads), suggesting that only a subset of supporting cells express TOX3, or that Sertoli cells are asynchronous in the timing of their expression of the protein. Interestingly, TOX3 positive AMH/SOX9 negative cells were also detected (Fig. 1G, yellow arrowheads).

### TOX3 knock down results in increased Leydig cell differentiation

To evaluate the role of TOX3 on testicular differentiation two different *TOX3* shRNAs (*sh370* and *sh685*) were cloned into *RCASBP(A)* viral vector expressing BFP reporter and the shRNA (37). To test the ability of the shRNA to knock down *TOX3* expression, DF-1 cells were transfected with RCASBP plasmids expressing one or other of these shRNAs. As a control, DF-1 cells were transfected with a non-silencing *shRNA* (37). After all cells had become BFP positive, they were transfected with *RCASBP(D)-GFP-T2A-TOX3* over-expression plasmid. 48 hours post-transfection, cells were fixed and *TOX3* knock down was assessed by reduction of GFP expression. *TOX3 sh685* showed the greatest reduction of GFP (and hence TOX3) expression (Extended data 1A). To quantify this reduction, flasks containing DF-1 cells were transfected with either *TOX3 sh685* or *non-silencing shRNA*. 72 hours post transfection, DF-1 cells were collected and plated in a 24 well plate and were left resting for 24 hours. Subsequently, cells were transfected with *RCASBP(D)-GFP-T2A-TOX3* over-expression plasmid and 48 hours post transfection they were collected for RNA extraction. *TOX3* expression was quantified by qRT-PCR, showing significant reduction (60%) of *TOX3* expression in cells expressing *TOX3 sh685*, compared with the control (Extended data 1B).

To evaluate the effect of knocking down *TOX3 in vivo*, RCASBP(A)TOX3-Sh685 virus or non-silencing control were injected into embryos at the blastoderm stage (day 0 of incubation). 100% of TOX3 knockdown embryos died at early stages of development, suggesting embryonic lethality when TOX3 knockdown occurred globally. To overcome this problem, *RCASBP(A)TOX3-Sh685* or non-silencing control plasmids were electroporated into the left coelomic epithelium of E2.5 chicken embryos. This afforded a more targeted delivery. Urogenital systems were harvested at E9.5 (stage 35/36), and immunofluorescence was performed on male (ZZ) gonads. As electroporation is innately variable, even across one gonad, we rely on localized immunofluorescence intensity as a measure of knockdown. While not quantitative, it is clear from Figure 2A that endogenous gonadal expression of TOX3 is lower after treatment with the specific shRNA. In the non-silencing control, TOX3 expression was uniform along the testicular cords, colocalizing with GFP (Fig. 2A). SOX9 expression was also reduced upon TOX3 knock down (Fig. 2B), whereas AMH or DMRT1 expression remained unchanged (Extended data 2A and B).

**Fig. 2.**
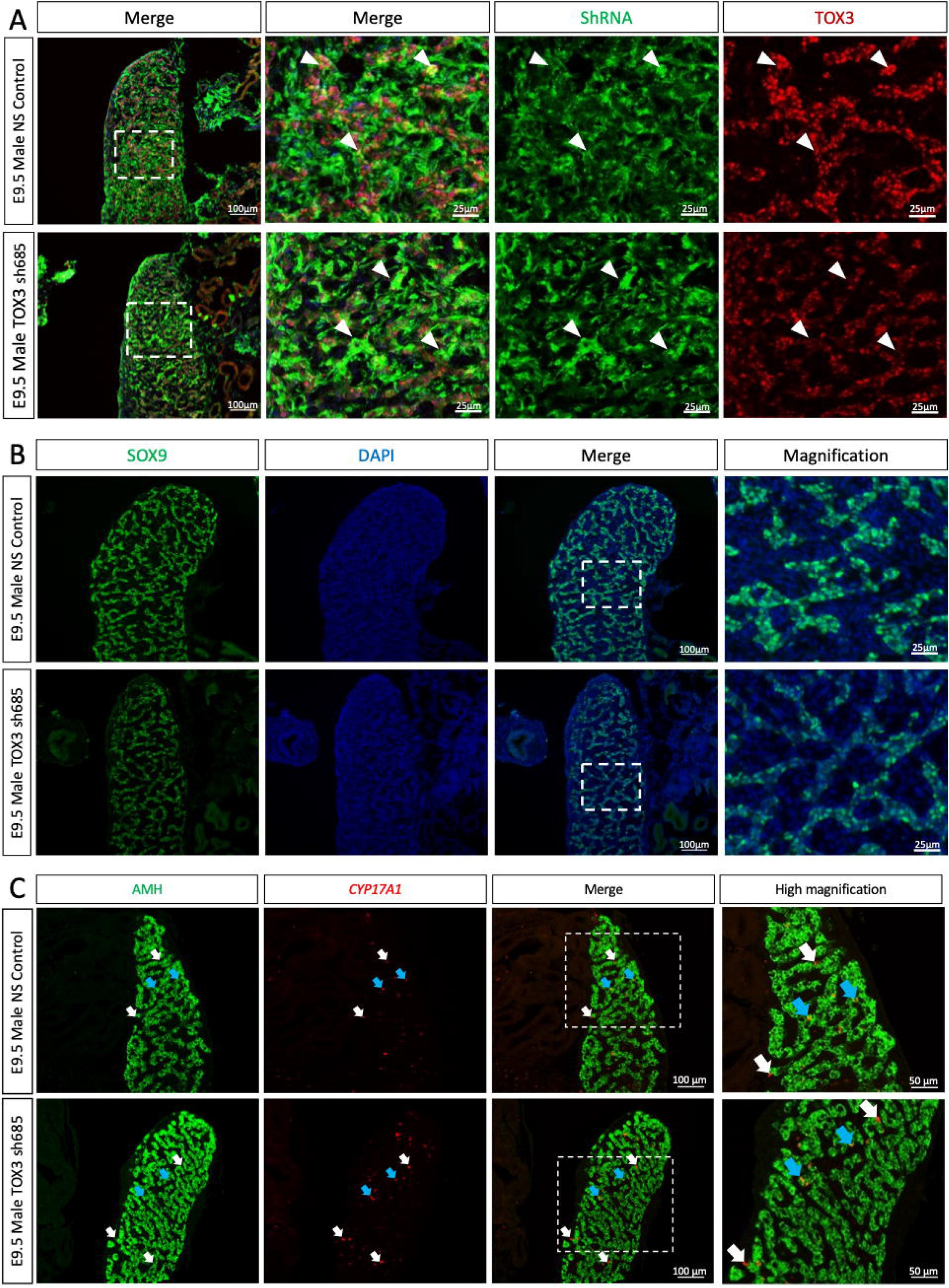
TOX3 knock down results in increased Leydig cell differentiation. *TOX3 sh685* or *NS shRNA* (control) plasmids were electroporated in chicken E2.5 coelomic epithelium. Immunofluorescence detection of (A) P27 (infection marker) and TOX3 in E9.5 male gonadal sections. White arrows indicate shRNA expressing supporting cells. (B) SOX9 immunofluorescence in control or TOX3 knock down E9.5 male gonads. (C) *CYP17A1* fluorescent in situ hybridization and AMH (Sertoli cell marker) immunofluorescence in TOX3 knock down or control E9.5 male gonads. Dashed box indicates the magnified area. White arrowheads indicate shRNA positive cells. White arrows indicate *CYP17A1* positive AMH negative Leydig cells. Blue arrows indicate intermediate cells (*CYP17A1* and AMH positive).

We previously reported that embryonic steroidogenic cells in the chicken embryo derive from a subset of previously committed supporting cells, in both males and females (23). This process of differentiation involves a sequential upregulation of the diagnostic steroidogenic cell marker *CYP17A1*, and a subsequent downregulation of the supporting cell markers (23). Our single-cell data shows that *TOX3* was not only expressed in the supporting cells, but also in a subpopulation of Sertoli cells that clusters transcriptionally with steroidogenic Leydig cells (Extended data 2C). Interestingly, *TOX3* is not expressed in the cells with the highest expression of *CYP17A1* (Extended data 2D), suggesting that *TOX3* might have a role in the Leydig cell differentiation. To address the role of *TOX3* in fetal Leydig cell differentiation, RCASBP(A)TOX3-Sh685 or non-silencing control plasmids were electroporated in the coelomic epithelium of E2.5 chicken embryos. Urogenital systems were harvested at E9.5, and *CYP17A1* fluorescent *in situ* hybridization was conducted to detect steroidogenic cells, followed by immunofluorescence against the Sertoli cell marker, AMH (Fig. 2C). *TOX3* knock down resulted in an increased number of *CYP17A1*+ Leydig cells, compared with the non-silencing control (Fig. 2C). This data suggests that TOX3 is required to inhibit Leydig cell differentiation.

### TOX3 over-expression in male gonads inhibits Leydig cell differentiation

To evaluate the effect of TOX3 in gonadal differentiation, *TOX3* open reading frame was cloned into RCASBP(D) viral vector (coupled to GFP reporter, as *RCASBP(D)-GFP-T2A-TOX3*). DF-1 cells were transfected with this construct and TOX3 protein and GFP expression were detected by immunofluorescence (Extended data 3A). Both nuclear TOX3 and cytoplasmic GFP were co-expressed in the transfected cells, validating the over-expression construct (Extended data 3A). Quantitative RT-PCR was also used to confirm *TOX3* mRNA over-expression in DF-1 cells, showing a significant increased expression, compared to the control (vector only expressing GFP) (Extended data 3B).

To address the role of *TOX3* in gonadal development *in vivo*, TOX3 was over-expressed in embryonic chicken gonads by coelomic electroporation at E2.5 using the RCASBP vector described previously. As a control, a RCASBP plasmid expressing GFP was electroporated. TOX3 was successfully over-expressed in E9.5 male gonads, colocalizing with GFP expression in the developing testis (Fig. 3A). No changes in Sertoli cell markers AMH, SOX9 and DMRT1 were detected when TOX3 was over-expressed in male gonads (Extended data 4A-C). Additionally, pre-granulosa marker aromatase was not detected in male gonads upon TOX3 overexpression (Extended data 4D). This suggests that TOX3 over-expression in male gonads has no effect in Sertoli cell differentiation.

**Fig. 3.**
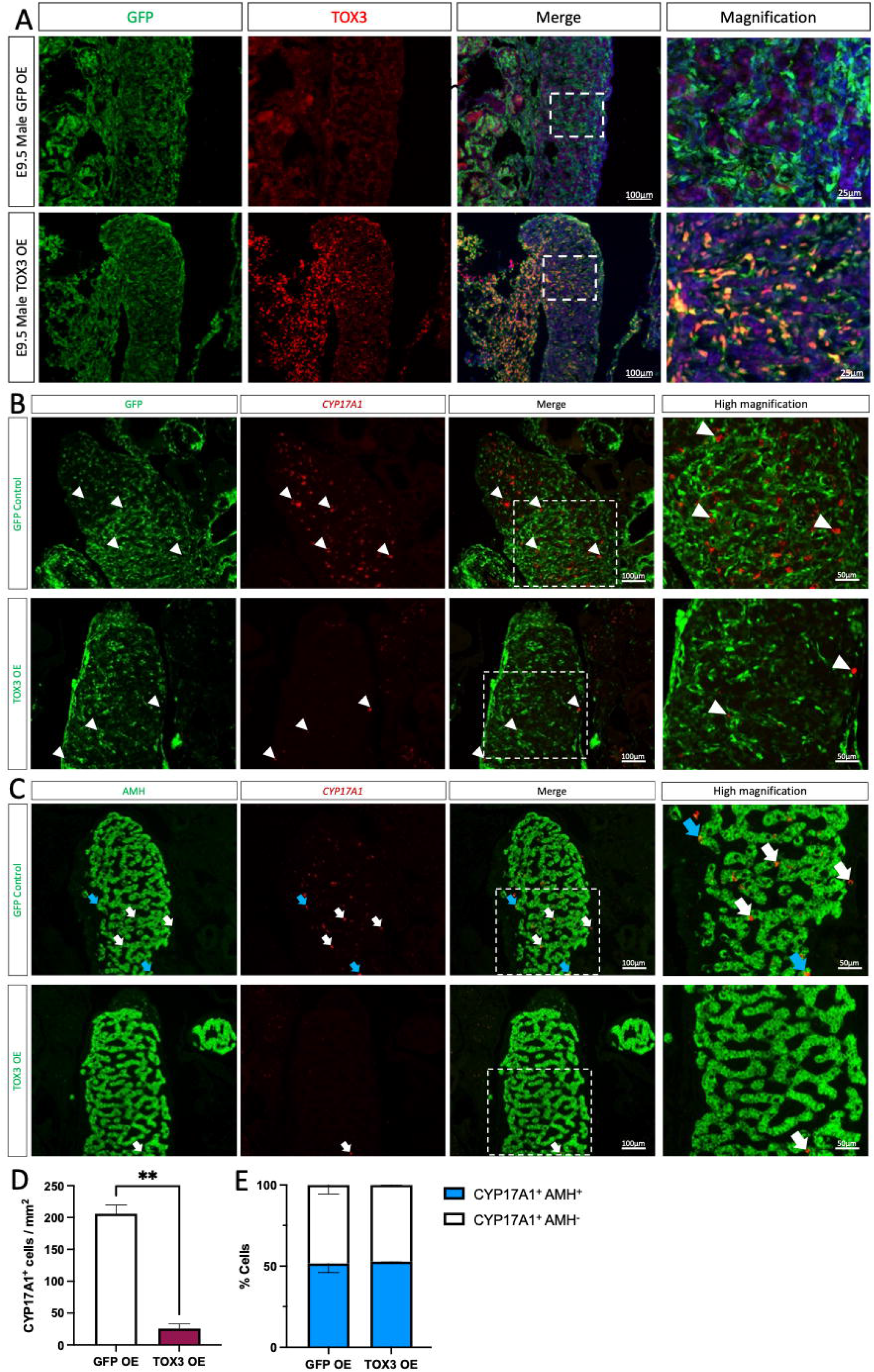
TOX3 over-expression in male gonads inhibits Leydig cell differentiation. TOX3 or GFP (control) over-expression plasmids were electroporated in chicken E2.5 coelomic epithelium. Male gonads were examined at E9.5. (A) immunofluorescence was performed to detect GFP and TOX3. (B) CYP17A1 fluorescent in situ hybridization was performed, followed by GFP immunofluorescence. (C) CYP17A1 fluorescent in situ hybridization was performed, followed by AMH immunofluorescence. Dashed box indicates the magnified area. White arrowheads indicate steroidogenic (CYP17A1 positive) cells. White arrows indicate CYP17A1 positive AMH negative Leydig cells. Blue arrows indicate intermediate cells (CYP17A1 and AMH positive). (D) Quantification of CYP17A1 positive cells per gonadal area in control (GFP OE) or TOX3 OE testis. Bars represent Mean ± SEM, n=3, **: p<0.01, unpaired two-tailed t-test. (E) Proportion of steroidogenic (CYP17A1+ AMH-) and intermediate (CYP17A1+ AMH+) cells in control (GFP OE) or TOX3 OE testis. Bars represent Mean ± SEM, n=3. 2-way ANOVA, Tukey’s post-test.

To evaluate the role of TOX3 overexpression in Leydig cell differentiation, *CYP17A1* fluorescent *in situ* hybridization was performed (Fig. 3B-C). Male gonads overexpressing *TOX3* showed 87% fewer *CYP17A1* positive cells than the control (Fig. 3B and D). Consistent with our previous finding that steroidogenic cells arise from a sub-population of AMH+ supporting (Sertoli) cells, CYP17A1+ cells were located both within the developing testis cords and outside of them. Interestingly, the ratios of “intermediate cells” (expressing both CYP17A1 and AMH) and Leydig cells (CYP17A1^+^ but AMH^-^) remained unaffected after TOX3 knockdown (Fig. 3C and E). Taken together, this data suggests that *TOX3* has a role in maintaining the identity of supporting cells by inhibiting the differentiation of steroidogenic Leydig cells and the induction of *CYP17A1* expression.

### TOX3 mis-expression in female gonads inhibits ovarian steroidogenic cell differentiation

To evaluate the role of *TOX3* in the ovarian differentiation, the gene was ectopically expressed by electroporation. *TOX3* was successfully mis-expressed in female gonads, colocalizing with GFP reporter expression in the developing ovary (Fig. 4A). Female gonads over-expressing TOX3 showed a reduction in aromatase expression in the region of the gonad mis-expressing TOX3, compared with the GFP control (Fig. 4B). The cortical region of the gonads was also affected, showing a thinner cortical structure in TOX3 mis-expressing ovaries (Extended data 5A). As cortical development is sensitive to estrogens (38), this could be a secondary effect of aromatase down-regulation. Despite reduction in the cortical domain in TOX3 expressing gonads, germ cells exhibited a normal localization in the cortical (non-medullary) region (Extended data 5B). Additionally, TOX3 over-expressing ovaries showed an increased expression of the male marker, AMH (colocalizing with GFP), compared with the control (Fig. 4C). No SOX9 or DMRT1 up-regulation was detected (Extended data 5C and D). In fact, DMRT1 appeared to be downregulated in female cells expressing TOX3 in the medulla (Extended data 5D).

**Fig. 4.**
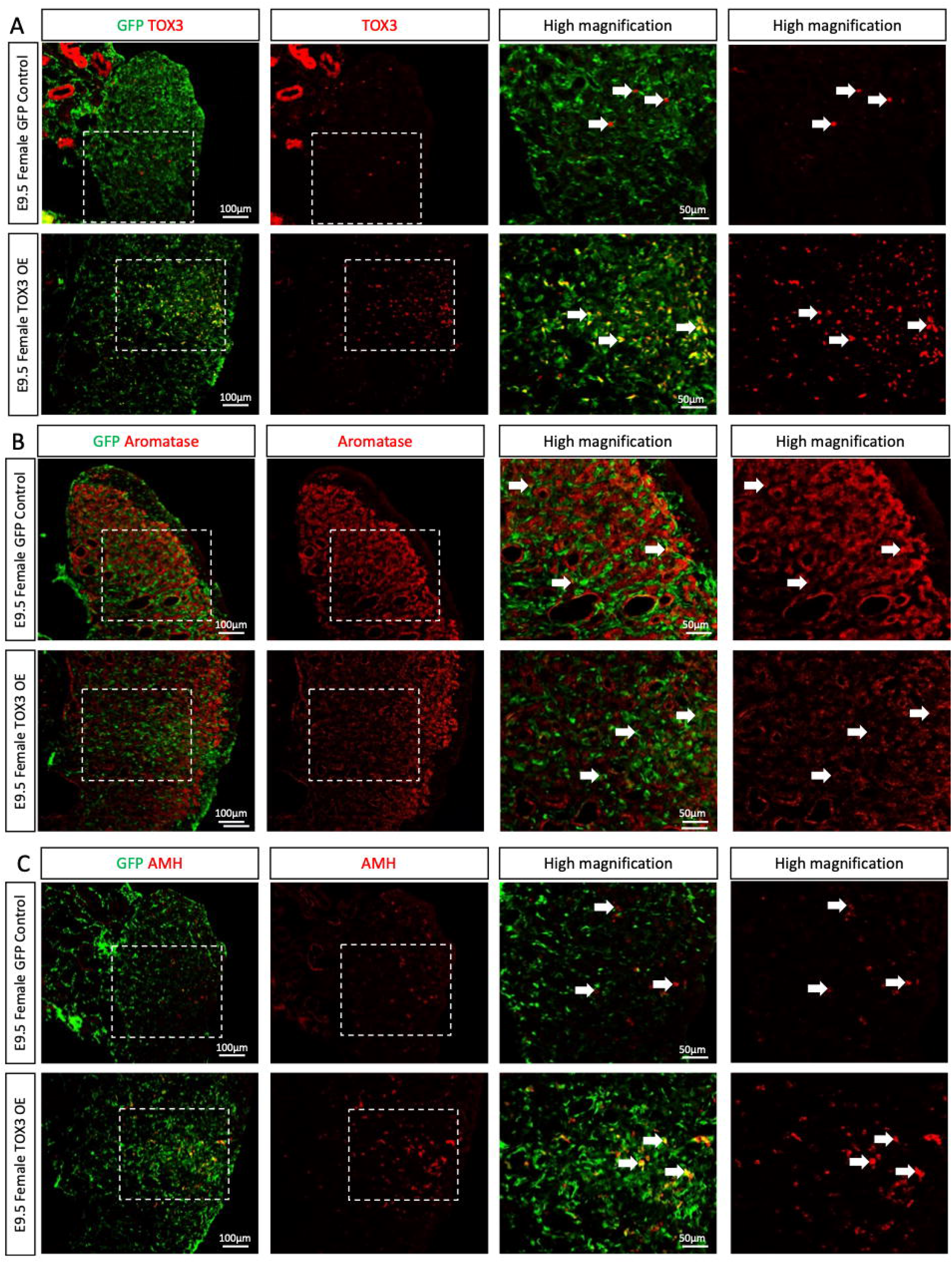
TOX3 over-expression in ovaries down-regulates aromatase. TOX3 or GFP (control) overexpression plasmids were electroporated in chicken E2.5 coelomic epithelium. Female gonads were collected at E9.5 and immunofluorescence was performed to detect GFP and (A) TOX3 (B), aromatase or (C) AMH. Dashed box indicates the magnified area. White arrow indicates GFP positive cells.

As chicken embryonic theca and Leydig cells share a similar transcriptome, with no evident sex specific markers (39), we tested if TOX3 mis-expression in females could modulate steroidogenic cell differentiation. Ovarian TOX3 mis-expression resulted in a reduction in the population of steroidogenic *CYP17A1* positive theca cells (Fig. 5A-C). In TOX3 mis-expressing gonads, GFP and CYP17A1 did not colocalize. This suggest that the remaining *CYP17A1* positive cells were TOX3 (GFP) negative (Fig. 5A). Additionally, TOX3 mis-expression resulted in AMH upregulation in females (Fig. 5B) concomitant with aromatase downregulation (Fig. 5C), consistent with the previous results. Taking together, the data suggests that, in females, TOX3 protein can inhibit estrogenic (CYP19A1+) and androgenic (*CYP17A1*+) cell differentiation.

**Fig. 5.**
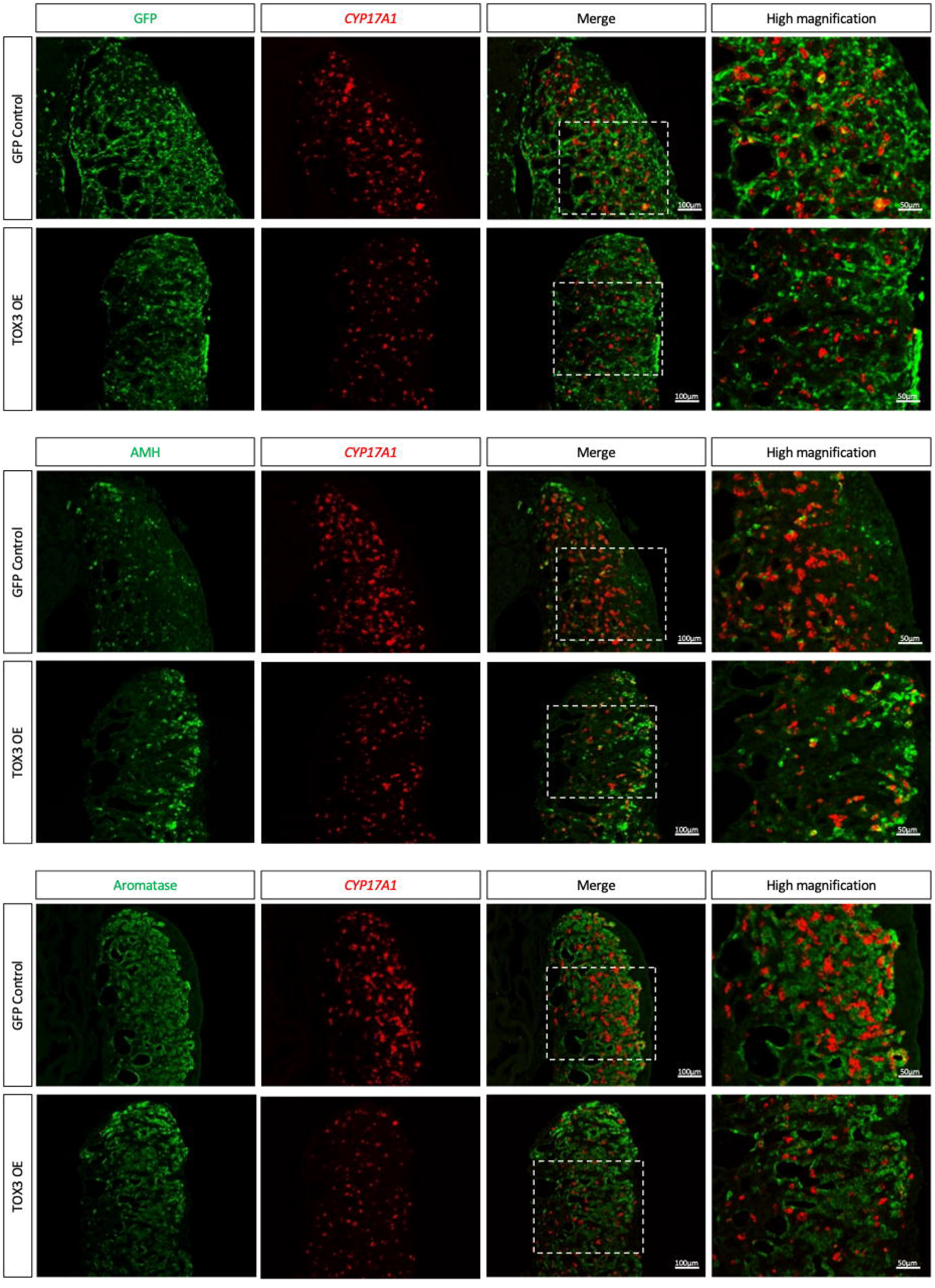
TOX3 overexpression inhibits steroidogenic theca cell differentiation. TOX3 or GFP (control) overexpression plasmids were electroporated in chicken E2.5 coelomic epithelium. Female gonads were collected at E9.5 and CYP17A1 fluorescent in situ hybridization was performed, followed by (A) GFP, (B) AMH or (C) Aromatase immunofluorescence. White arrowheads indicate steroidogenic (CYP17A1 positive) cells. Dashed box indicates the magnified area.

### Estrogen inhibits TOX3 expression

Steroid hormones influence avian gonadal sex differentiation. Specifically, estrogen is required for ovarian differentiation (40). The estrogen synthesizing enzyme, aromatase, is activated only in female gonads at the onset of gonadal sex differentiation, and the estrogen that it produces has a positive feedback effect upon further aromatase gene transcription. Estrogen synthesized in the gonadal medulla regulates the adjacent cortex, inducing proliferation (38). Conversely, exposure of male embryo to exogenous estrogens can feminize gonads (38, 41-44). To understand the regulation of *TOX3* expression, 17β-estradiol (E2) or vehicle (Oil) were injected into chicken eggs at E3.5, prior to the onset of gonadal sex differentiation and male up-regulation of TOX3. E9.5 urogenital systems were then collected and processed for *aromatase, AMH* and *TOX3* whole mount *in situ* hybridization (Fig. 6A and B) or for qRT-PCR (Fig. 6C).

**Fig. 6.**
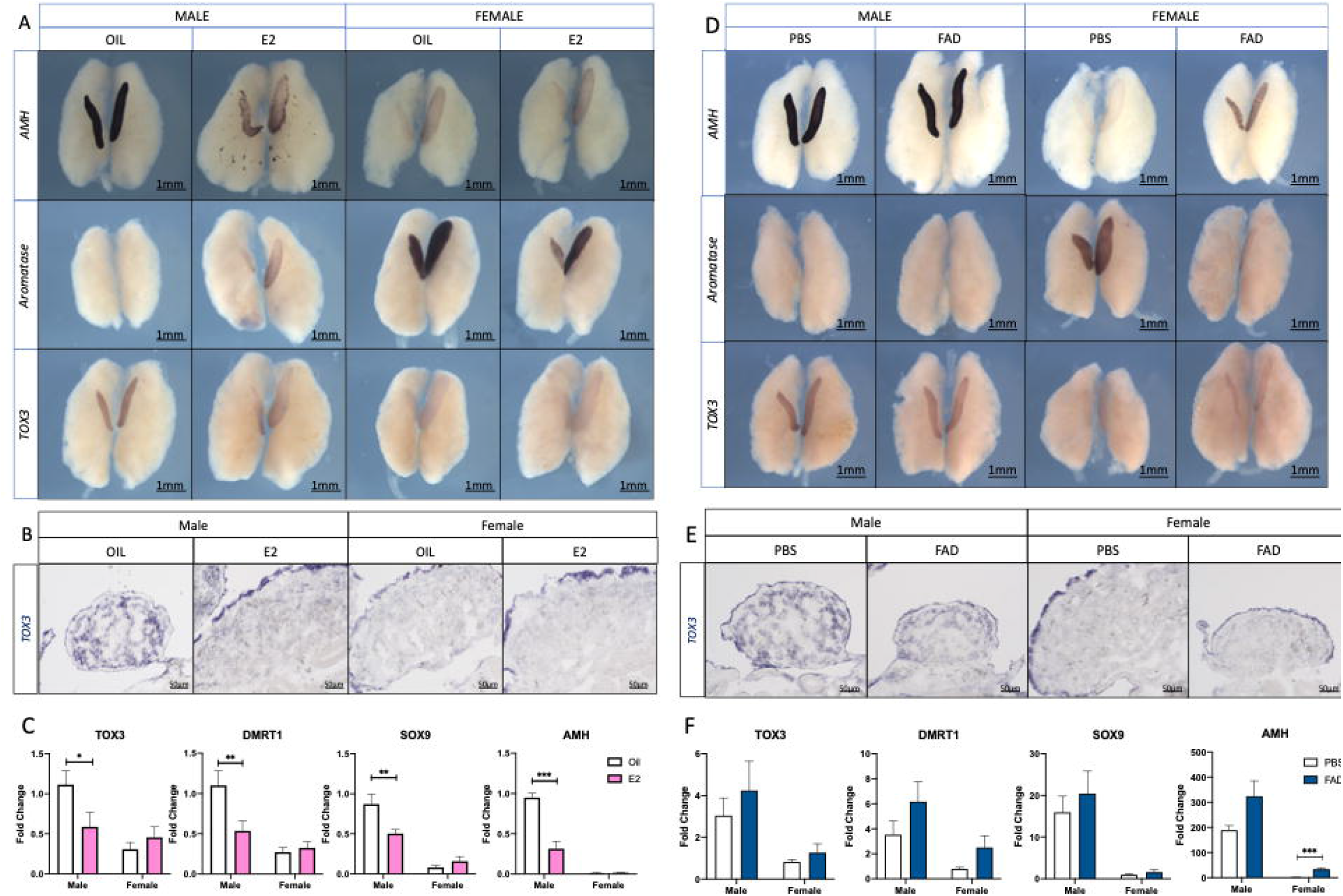
*Estrogen inhibits TOX3 expression*. (A) 17β-estradiol (E2) or vehicle (Oil) were injected in E3.5 chicken eggs. Male and female gonads were collected at E9.5 for TOX3 whole mount *in situ* hybridization (A and B) or for qRT-PCR (C). (A) Down-regulation of *AMH*, up-regulation of *aromatase* and down-regulation of *TOX3* in male gonads exposed to E2 compared to Oil control. (B) *TOX3* WISH overstained sections. (C) *TOX3, DMRT1, SOX9* and *AMH* mRNA expression levels by qRT-PCR. Expression is relative to β-actin and normalized to Male Oil. (D) Fadrozole (FAD) or vehicle control (PBS) were injected in E3.5 chicken eggs. Male and female gonads were collected at E9.5 for TOX3 WISH (D and E) or qRT-PCR (F). (D) Up-regulation of AMH, down-regulation of aromatase but no effect upon TOX3 mRNA expression in female gonads. (E) TOX3 WISH overstained sections. (F) qRT-PCR for TOX3, DMRT1, SOX9 and AMH. Expression is relative to β-actin and normalized to Female PBS. Bars represent Mean ± SEM. *,**,*** = p<0.05, 0.01 and 0.001 respectively. Multiple t-test and Holm-Sidak post-test.

Male left gonads exposed to estrogens were larger than the controls, showing a female-like left-right asymmetry and suggesting male-to-female sex reversal. *AMH* and *TOX3* expression were reduced in male gonads exposed to 17β-estradiol (E2) (Fig. 6A), whereas expression of the normally female-restricted marker, aromatase, was induced in males (Fig. 6A). Gonadal sections of *TOX3* WISH revealed that male left gonads exposed to E2 had an ovarian like structure, which lacked *TOX3* expression in the medulla (Fig. 6B). *TOX3* expression levels were quantified by qRT-PCR, showing a significant reduction of the expression level in male gonads exposed to 17β-estradiol (E2), compared with the control (Oil) (Fig. 6C). Additionally, Sertoli cell markers *SOX9, DMRT1* and AMH also showed a significant expression reduction (Fig. 6C).

Inhibiting aromatase function with the drug fadrozole induces masculinization of female gonads (43, 45, 46). When female embryos were treated with fadrozole, *TOX3* was not upregulated in the gonad (Fig. 6D-F). This was despite the down-regulation of aromatase (Fig. 6D). Furthermore, *AMH* but not *SOX9* nor *DMRT1* were significantly up-regulated in feminization experiments (Fig. 6D and F). Taken together, this data suggests that *TOX3* expression is negatively regulated by estrogens, directly or indirectly. However, the absence of estrogens is not sufficient to induce *TOX3* expression, pointing to the requirement of a male-specific factor.

### DMRT1 regulates *TOX3* expression

The Z-linked transcription factor, DMRT1, is the master testis-determinant in chicken. *DMRT1* knock out or knock down induces feminization or complete ovary formation (12, 44, 47). Targeted over-expression of *DMRT1* alone induces upregulation of *SOX9* and *AMH* (male specific genes) in female chicken gonads (19). In mammals, the main role of the sex determining gene *Sry* is to induce the expression of *Sox9*, which promotes testicular differentiation by activating pro-testis genes and repressing pro-ovarian genes (6, 48-50). To understand how *TOX3* is regulated in the embryonic chicken testis, DMRT1, SOX9 or GFP (as a control) were over-expressed in DF-1 cells, a chicken fibroblastic cell line. *DMRT1, SOX9* and *TOX3* mRNA expression levels were evaluated by qRT-PCR. DMRT1 over-expression, but not SOX9, was able to significantly upregulate *TOX3* expression *in vitro* (Fig. 7A).

**Fig. 7.**
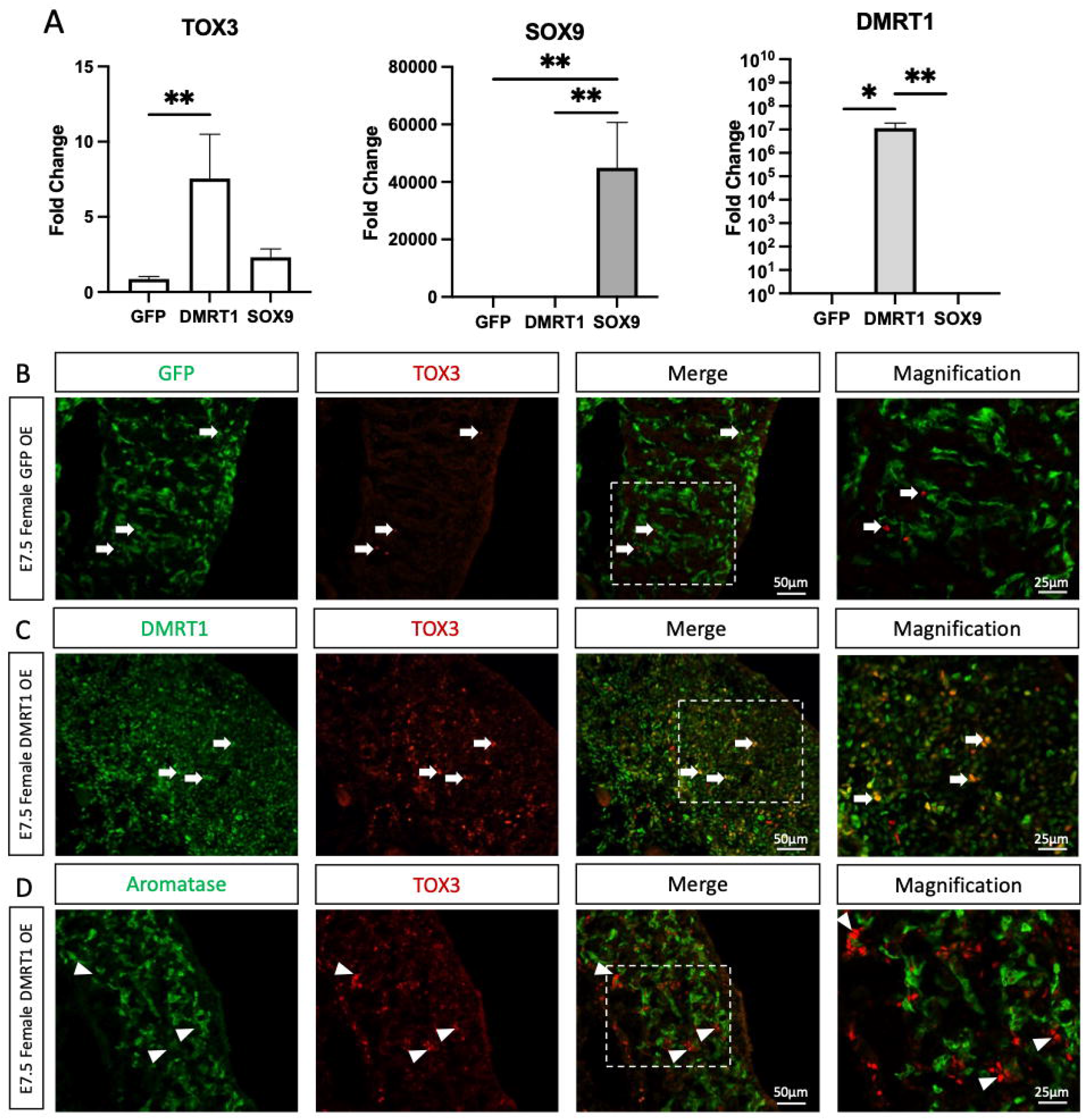
DMRT1 overexpression induces TOX3 expression *in vitro* and *in vivo*. (A) DMRT1, SOX9 or GFP as a control were overexpressed *in vitr*o in DF-1 chicken fibroblastic cells and *TOX3, SOX9* and *DMRT1* mRNA expression was measured by qRT-PCR. Expression is relative to *β-actin* and normalized to GFP overexpression control. Bars represent Mean ± SEM. *,** = p<0.05 and 0.01 respectively. Multiple t-test and Holm-Sidak post-test. (B)TOX3 and GFP, (B) DMRT1 or (C) aromatase immunofluorescence in E7.5 female gonadal sections overexpressing GFP (control) or DMRT1. Dashed box indicates the magnified area. White arrows indicate TOX3 positive cells. White arrowheads indicate TOX3 positive Aromatase negative cells.

To evaluate if it was sufficient to induce *TOX3* expression *in vivo, DMRT1* was over-expressed in chicken gonads by coelomic electroporation at E2.5, as reported previously (19). As a control, a GFP expressing plasmid was electroporated. Immunofluorescence for TOX3 was performed on E7.5 female gonads. In female control gonads, TOX3 protein expression was minimal (Fig. 7B). In contrast, when DMRT1 was mis-expressed in female gonads, TOX3 was upregulated, colocalizing with DMRT1 (Fig. 7C). In addition, TOX3 positive cells did not colocalize with the pre-granulosa marker aromatase, suggesting that TOX3 is in fact expressed in DMRT1-induced Sertoli cells (Fig. 7D). To evaluate if it is necessary to regulate *TOX3* expression, *DMRT1* was knocked down by viral injection, using the blastoderm delivery of an shRNA method reported previously (12). In these experiments, a GFP reporter was used, marking those cells infected with virus expressing the DMRT1-specific shRNA or the control (scrambled shRNA). The *DMRT1* shRNA has been previously validated and published by our laboratory, showing robust knockdown of *DMRT1* mRNA and protein expression (12). Control E9.5 male gonads expressing the non-silencing control shRNA (scrambled shRNA) showed normal DMRT1, SOX9, AMH and TOX3 expression in the testicular cords, and no expression of female pre-granulosa markers (FOXL2 and aromatase) (Fig. 8). In contrast, aromatase and FOXL2 were locally up-regulated in male gonads in regions where DMRT1 expression was knocked down, colocalizing with the GFP reporter (and hence shRNA delivery) (Fig. 8). GFP positive cells were TOX3 negative, indicating that in the cells where DMRT1 was down-regulated, TOX3 expression was not expressed, or lost (Fig. 8). Other Sertoli markers, SOX9 and AMH were also absent in the DMRT1-knockdown region of the gonad, not colocalizing with GFP (Fig. 8), consistent with our previous data (12). Taken together, these data indicate that DMRT1 is necessary and sufficient to induce *TOX3* expression.

**Fig. 8.**
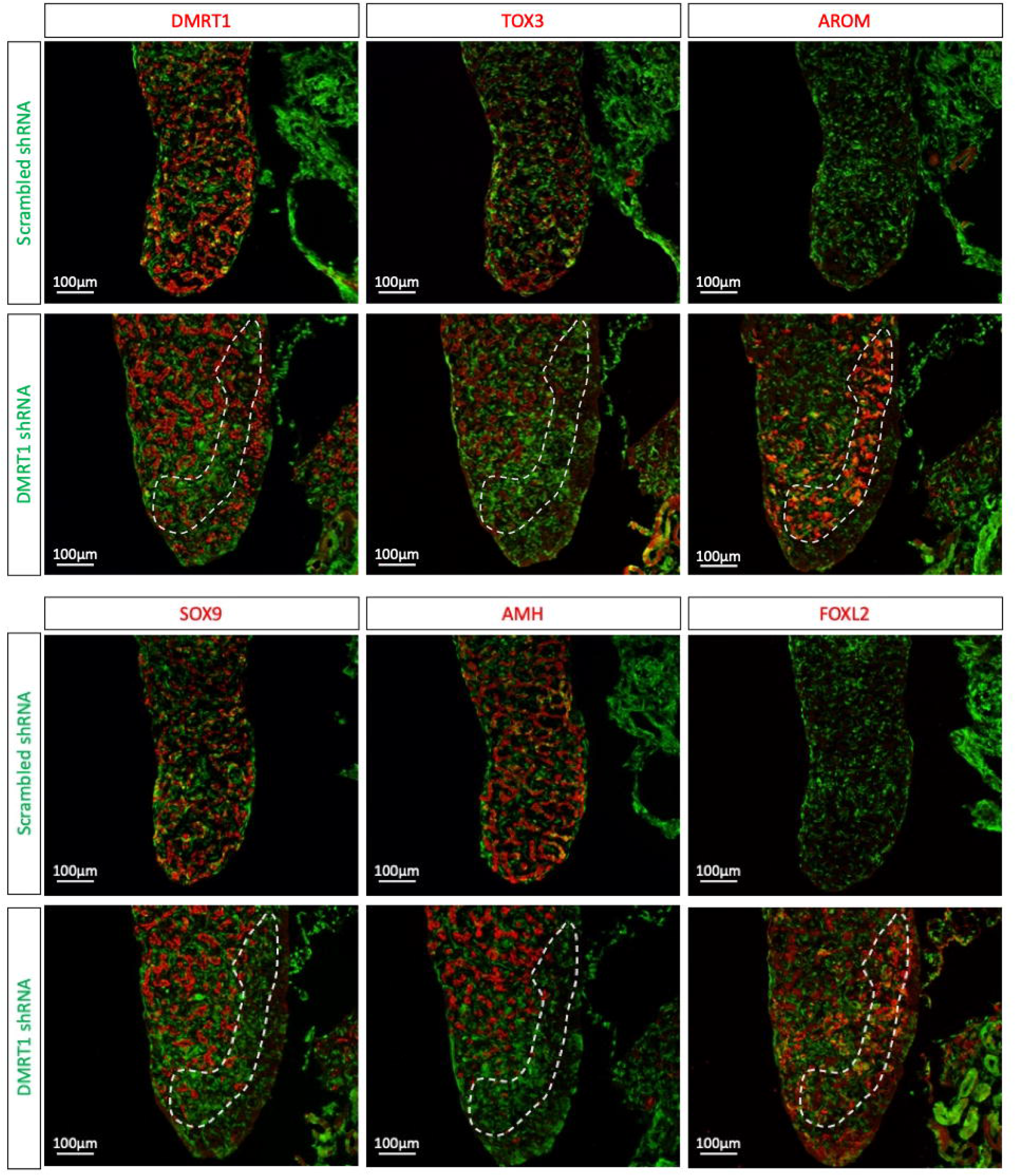
*In vivo* DMRT1 knock down inhibits TOX3 expression in male gonads. RCAS virus expressing *DMRT1 shRNA343* or *scrambled shRNA* were injected at blastoderm stage. Gonads were examined at E9.5 and immunofluorescence against GFP (transfection marker), DMRT1, TOX3, aromatase (AROM), SOX9, AMH and FOXL2 was performed. Dashed area indicates a DMRT1 knock down gonadal region in supporting cells.

## Discussion

During gonadal sex differentiation, gene regulatory networks control commitment of the supporting cell lineage into Sertoli cells in the testis and pre-granulosa cells in the ovary (51). In mammals, *Sry* is the molecular trigger for the Sertoli cell differentiation program (4, 52, 53). *Sry* is absent in birds and a different sex chromosome system applies. Avians have a ZZ male; ZW female sex chromosome system, un-related to the X and Y of mammals. In the most widely studied avian model, the chicken, Sertoli cell differentiation is directed by the Z chromosome-linked gene *DMRT1* (44, 47). In the chicken embryo, *DMRT1* expression is confined to the urogenital system. It is expressed in the supporting cell progenitors of both sexes (54). However, due to the lack of global Z chromosome dosage compensation in birds, *DMRT1* is expressed at approximately twice the level in male gonads compared to female gonads (55-57). This higher level of expression is sufficient to induce the testicular developmental program in ZZ embryos (19, 56). Consistent with this hypothesis, *DMRT1* knockdown and single allele knock out in males result in pre-granulosa cell differentiation and ovary formation (12, 47). Conversely, DMRT1 over-expression in female (ZW) gonads results in Sertoli cell differentiation and testicular development (19). Despite its key role in chicken Sertoli cell differentiation, the exact mechanism of action and DMRT1 direct targets in the chicken gonad are not completely known (58).

In this study, we showed that the HMG-box transcription factor gene, *TOX3*, has a novel role in chicken gonadal sex differentiation, acting downstream of *DMRT1*. Firstly, TOX3 is expressed in a sexually dimorphic manner during gonadal development, being male up-regulated. The gene is down-regulated in male gonads exposed to exogenous estrogen, though not up-regulated in female gonads with experimentally disrupted estrogen synthesis. Knockdown of TOX3 resulted in a lower level of SOX9 expression in male gonads, but testis cords still formed and AMH was still expressed (Fig 2). However, the steroidogenic population, as assed by CYP17A1, was enhanced. Conversely, over-expression of TOX3 resulted in blunted steroidogenic cells, in both sexes. These observations indicate the TOX3 has a role in modulating steroid cell differentiation, acting downstream of DMRT1. *DMRT1* knockdown in male gonads results in localized loss of TOX3 protein expression, while DMRT1 mis-expression in females causes up-regulation of TOX3 expression. Over-expression of TOX3 in female gonads disrupts proper aromatase expression and activates AMH. Conversely, TOX3 knockdown resulted in downregulation of SOX9, whereas DMRT1 and AMH expression remained unchanged. Over-expression in both male and female gonads reduced the pool of CYP17A1+ steroidogenic cells, whereas knockdown in males resulted in an increment in the number of CYP17A+ cells.

We first identified *TOX3* as a novel gene expressed in pre-Sertoli cells based on a bulk RNA-seq screen (23). *TOX3* mRNA was found to be up-regulated in male but not female gonads after the onset of gonadal sex differentiation (between E6.5/stage 30 and E8.5/stage 35). TOX3 protein was not detected at E6.5, suggesting that there is a translational delay or that the levels of protein expression were not high enough to be detected by immunostaining. Additionally, TOX3 protein expression was not homogeneous in the testicular cords, colocalizing partially with AMH and SOX9. This could reflect expression in Sertoli cells at different (asynchronous) stages of development or could reflect expression in a distinct subset of cells. These findings agree with the chicken testicular single-cell RNA-seq where two different Sertoli cell types were identified, suggesting that the Sertoli cell population is not as homogeneous as generally considered (23). A similar phenomenon has also been described for the chicken Z-linked gene *HEMGN*, which exhibits variable levels of expression in the nuclei of developing male gonads (19). *HEMGN* encodes a transcription factor expressed in developing chicken Sertoli cells and required for testis development (59). It lies downstream of DMRT1 in the male developmental pathway in chicken, though it might not be conserved in other birds(44). It would be of interest to determine whether the variable TOX3 expression spatially coincides with the HEMGN^+^ Sertoli cells.

The data presented here indicate that DMRT1 activates *TOX3* during testicular development in the chicken embryo. *TOX3* is likely to be one of many genes activated by DMRT1, generating a network of factors coordinating testis formation and function. We and others have previously shown that DMRT1 directly or indirectly regulates *SOX9* and *AMH* during chicken testis development (19, 47). We hypothesize that DMRT1 directly regulates *TOX3* and *SOX9* expression. Mis-expression of TOX3 in female gonads had an effect upon the endocrine development of the gonad, by locally activating AMH and suppressing aromatase expression. As *TOX3* knockdown in males did not affect AMH expression, it is likely that ectopic TOX3 in the female gonad affects AMH expression indirectly by suppressing aromatase expression. Estrogen and AMH are mutually antagonistic in the avian gonad (18, 20, 46). Suppression of aromatase, and hence estrogen synthesis, allows up-regulation of *AMH* expression in the female gonad (Fig. 6D) (60). Conversely, exogenous estrogen (E2, 17-β-estradiol) causes AMH mRNA down-regulation in male gonads (Fig. 6A). Hence, here, TOX3 may have led to increased AMH in the female chicken gonad via local suppression of aromatase. In the male gonad, one of the roles of TOX3 in the supporting cell lineage (Sertoli cells) may be to ensure that the aromatase gene is inactive, hence preventing feminization. It would be of interest to scrutinize the *CYP19A1* gene, which encodes aromatase, for TOX3 binding sites.

In the female avian embryo, ovarian cortex development is dependent on estrogens, which are synthesized in the medullary cord cells (pre-granulosa type cells). Accordingly, ERα is expressed in the gonadal cortex (18, 38). When *TOX3* was over-expressed in female gonads, the cortical area was reduced. This could also be a secondary effect of the reduction in aromatase expression, the enzyme responsible for transforming androgens into estrogens. Additionally, TOX3 mis-expression inhibited steroidogenic embryonic theca cell differentiation (Fig. 5). Fewer embryonic theca cells may cause a reduction in androgens and, as androgens are the substrate for estrogen synthesis, in lower levels of estrogens and thus a reduced cortex. Surprisingly, despite having a thinner cortex, germ cells remained in the cortical or juxtacortical region and not in the medulla when TOX3 was mis-expressed in females. This could suggest that in fact, the germ cells are not surrounded by epithelial cells but from so-called juxtacortical medulla cells. This could explain previous reports of FOXL2 positive cells, presumably pre-granulosa cells, present in the juxtacortical medullary region of E14.5 chicken ovaries (17).

The data presented here indicates that another role of *TOX3* in the developing chicken gonad is modulation of steroidogenic cell differentiation. In the embryonic chicken testis, we have shown that the *CYP17A1*^+^ pre-Leydig cell progenitors likely derive from a subset of the Sertoli cells (23). When TOX3 over-expression was performed, there was a clear loss of *CYP17A1* expression, presumably reflecting a loss of steroidogenic progenitor cells. This suggests that TOX3 may normally act to restrain or modulate the sub-population of Sertoli cells that differentiate into the *CYP17A1*^+^ embryonic steroidogenic Leydig cells around the seminiferous cords. During this Sertoli to Leydig cell differentiation process, a subset of AMH positive cells upregulate steroidogenic markers, such as *CYP17A1* (23). We consider these to be the transitioning population (Sertoli to Leydig cells). It is unknown if all the Sertoli cells have the ability to differentiate into steroidogenic cells or if specific factors are required to control this process. *TOX3* appears to be a key factor in this cell fate decision. As noted, not all AMH positive pre-Sertoli cells were also TOX3 positive. Additionally, TOX3 over-expression in male gonads resulted in a reduction of both steroidogenic cells (*CYP17A1*^*+*^) and the intermediate/transitioning cells (AMH^+^/CYP17A1^+^). This suggests that *TOX3* inhibits or modulates the Sertoli to Leydig cell transition and maintains Sertoli cell identity. Based on this concept, TOX3^-^/AMH^+^ cells may differentiate into Leydig cells whereas TOX3^+^/AMH^+^ cells may follow the Sertoli cell differentiation path. Further experiments involving lineage tracing and fate mapping are required to validate this theory. It is interesting, in this regard, that TOX3 down-regulation has been reported to facilitate epithelial to mesenchymal transition by repression of *SNAI1* and *SNAI2* in cancer cells (30). This would be consistent with our findings, as TOX3 over-expression appears to suppress the proposed transition of Sertoli (epithelial) to Leydig (mesenchymal) cells, as based on *CYP17A1* expression.

Alternatively, TOX3 expression in the supporting cell lineage may act indirectly by inducing paracrine factors that are secreted to regulate differentiation of the steroidogenic Leydig lineage. However, there are some caveats to these interpretations. Firstly, the expression of *CYP17A1* is taken as a marker of the steroidogenic population. Loss of *CYP17A1* mRNA expression could reflect fewer pre-Leydig cells developing, as interpreted here, or it could simply reflect downregulation of that gene in the Leydig population. It would be of interest to examine *CYP17A1* expression together with cell proliferation markers and other markers of the steroidogenic lineage, such as *CYP11A1*, to distinguish these possibilities. In the mouse, 3β-HSD is widely used as a diagnostic marker of the male steroidogenic (fetal Leydig cell) population. However, in chicken, 3β-HSD is expressed in the embryonic Sertoli cells, not in the Leydig cell progenitors, and is therefore not an appropriate marker for the latter (20).

Human DNA variants in the *TOX3* locus has been associated with polycystic ovarian syndrome (PCOS) in several populations (31-34). These mutational variants are not found in the coding region of *TOX3*, indicating that *TOX3* transcriptional regulation might be affected (61). PCOS is one of the most common infertility causes in females (62). Hyperandrogenism (elevated androgen) is one of the main diagnostical caracteristics of this syndrome (63). In adult PCOS granulosa cells, lower levels of TOX3 protein are detected, compared with the control (64). This suggests that lower levels of ovarian *TOX3* expression might act as causative factor for PCOS.

The exact functional mechanism of *TOX3* in this disease or in the ovarian context is not known (31). The data presented here demonstrate that *TOX3* inhibits the differentiation of steroidogenic cells in embryonic chicken gonads. In PCOS, lower levels of *TOX3* could be implicated in an increase in steroidogenic producing cells, explaining the higher levels of androgens described in PCOS ovaries (65). This is consistent with our *TOX3* knock down data (Fig. 2C). Additionally, rats with global lower levels for *Tox3* display obesity, sterility (male and female) and increased anxiety, all of which are symptoms of PCOS (65). Taking together this indicates that downregulation of *TOX3* in ovaries could lead to PCOS in females by a dysregulation of steroidogenic cell differentiation. It has been suggested that dysregulated or abnormal embryonic gonadal development could be the cause of PCOS (66). Several PCOS associated genes are expressed at different key developmental stages and in different cell types during ovarian development (66). In contrast, the expression levels of PCOS associated genes have not been studied in testicular development, an organ specialized in androgen production. The current study described the role of a testicular associated gene in polycystic ovarian syndrome, suggesting that abnormal embryonic gonadal sex differentiation could be one of the causes for PCOS. Additionally, further research is required to assess the expression levels and cell types where *TOX3* is expressed in adult gonads.

PCOS is a multifactorial disease, being associated with genetic and environmental factors (67). This syndrome shows a transgenerational inheritance, involving both genetic and epigenetic factors (68, 69). It is unclear if the high levels of androgens could be the cause or a consequence of PCOS. Animals treated with androgens for long time periods develop all the symptoms of PCOS, suggesting an important role in the pathogenesis of the disease (70-72). Interestingly, *TOX3* expression decreases in both the ovary and the brain in a high testosterone PCOS zebrafish model (72). This indicates that not only genetic mutations can cause the dysregulation of *TOX3* in PCOS patients, but that hormonal levels play a role in its regulation.

The data presented here indicate that the chicken embryo may serve as a potential model for PCOS. TOX3 modulates CYP17A1+ steroidogenic lineage development. Meanwhile, *TOX3* expression is repressed by estrogen (Fig. 6). Further research should focus on the role of androgens and estrogens in regulating *TOX3. TOX3* could have a dual role in PCOS pathogenesis. It would be interesting to examine the effect of exogenous androgen or androgen antagonists on TOX3 expression in the chicken model. While DMRT1 is necessary and sufficient for TOX3 expression in chicken, maintenance of its expression may involve an endocrine control. In humans, *TOX3* downregulation by genetic mutations in regulatory regions may result in steroidogenic cell proliferation and thus elevated androgen synthesis. On the other hand, higher levels of androgens, not associated with *TOX3*, can inhibit *TOX3* expression, which in turn results in more androgenic cells in the gonad. *TOX3* could be a new potential target for novel PCOS therapies.

The data presented describes the role of *TOX3* in potentially maintaining Sertoli cell identity and inhibiting the steroidogenic lineage in embryonic chicken gonads. However, the mechanism is unclear. Omics technologies will be crucial to uncover *TOX3* function in gonadal differentiation. *In ovo* overexpression and knockdown experiments can be coupled with RNA sequencing to identify genes regulated by *TOX3*. Additionally, TOX3 ChIP-seq will determine the direct target genes that TOX3 regulates. Furthermore, comparative analysis with other organisms is required to fully understand the role of *TOX3* in gonadal development and, potentially, PCOS progression.

## Methods

### Eggs & Sexing

HyLine Brown fertilized chicken eggs (*Gallus gallus domesticus*) were obtained from Research Poultry farm (Victoria, Australia) and incubated at 37°C under humid conditions. Embryos were staged *in ovo* according to Hamburger and Hamilton (36). Sexing was performed by PCR, as described previously (73). ZW females were identified by the presence of a female-specific (W-linked) *XhoI* repeat sequence in addition to a *18S* ribosomal gene internal control. ZZ males showed the 18S band only (73).

### Sex reversal experiments

Sex reversal experiments were performed as described previously (37). Briefly E3.5 (HH stage 19) eggs were injected with 1.0 mg of Fadrozole (Novartis), PBS (Vehicle), 0.1mg of E2 in 10% Ethanol in Sesame oil solution or vehicle (Control). Urogenital systems for WISH or gonads for qRT-PCR were collected at E9.5 (HH35).

### Whole mount *in situ* hybridization

Whole mount in situ hybridization was performed as described before (23). Briefly urogenital systems were fixed overnight in 4% PFA in DEPC-PBS. After methanol dehydration and rehydration to PBTX (PBS + 0.1% Triton X-1000), tissues were permeabilized in proteinase K 10 mg/mL for up to 2 hours. Tissues were briefly re-fixed and placed into pre-hybridization solution overnight at 65°C. *TOX3* (23), *AMH* (74) and *aromatase* (44) antisense probes were added to pre-hybridized tissues (approx. 7.5 µL/tube) and hybridization was carried out overnight at 65°C. Tissues were then subjected to stringency washes, blocked in TBTX/BSA/Sheep serum and then treated overnight with anti-DIG-AP antibody (1:2000; Roche). Following extensive washing in TBTX, tissues were exposed to BCIP/NBT color reaction at room temperature for up to 3 hours. Color reaction was stopped at the same time for each gene by rinsing in NTMT buffer, TBTX, PBTX, PBS and imaging. To examine tissue sections, samples were overstained for 2 days, cryoprotected in PBS plus 30% sucrose, snap frozen in OCT, and cryosectioned (10 µm).

### RNA extraction and qRT-PCR

Gonadal pairs were collected in 330 µl of Trizol reagent (ThermoFisher) and kept at -80°C until processing. After sexing, 3 same sex gonadal pairs were pooled for each sample, homogenized and followed RNA extraction method as per the manufacturer’s instructions (Trizol, ThermoFisher). For chicken DF-1 cells, confluent cells on each well (24 well plate) were collected in 1 ml of Trizol reagent (ThermoFisher) and stored at -80°C until processing. Genomic DNA was removed using DNA-free™ DNA Removal Kit (Invitrogen) and 500 ng-1µg of RNA was converted into cDNA using Promega Reverse Transcription System (A3500). RT-qPCR was performed using QuantiNova SYBR® Green PCR Kit. Expression levels were quantified by the 2^-ΔΔCt^ method using *β-actin* as housekeeping internal control gene. Data was analysed using unpaired t-test (two groups), multiple t-tests (one per embryonic stage/treatment) or one-way nonparametric ANOVA (if more than two groups were analysed). Statistical significance was determined using the Holm-Sidak method for the multiple t-test or Tukey test for ANOVA. Primers: *TOX3* Fw: TCAGAGCTTGGATCTCCCCT, *TOX3* Rv: *GGCGATACTGCGAAACTTGG, SOX9* Fw: *GTACCCGCATCTGCACAAC, SOX9* Rv: *TTCTCGCTCTCATTCAGCAG, DMRT1* Fw: *GGACTGCCAGTGCAAGAAGT, DMRT1* Rv: *GGTACAGGGTGGCTGATCC, AMH* Fw: *GAAGCATTTTGGGGACTGG, AMH* Rv: *GGGTGGTAGCAGAAGCTGAG, β-actin* Fw: *GCTACAGCTTCACCACCACA, β-actin* Rv: *TCTCCTGCTCGAAATCCAGT*.

### Immunofluorescence

Embryonic urogenital systems were briefly fixed in 4% PFA/PBS, cryo-protected in 30% sucrose, blocked into OCT, snap frozen and 10 µm frozen sections were then cut. Immunofluorescence was carried out as described before (23). Briefly, sections were permeabilized in 1% Triton X-100 in PBS for 10 min at room temperature, blocked in 2% BSA/PBS for 1hr at room temperature followed by primary antibody incubation overnight at 4°C. The following primary antibodies were used: goat anti-GFP antibody (Rockland, 1:500), mouse anti-pan cytokeratin (Novus Bio, 1:200), rabbit anti-DMRT1 (in house antibody; 1:2000), rabbit anti-SOX9 (Millipore antibody, 1:4000), rabbit anti-AMH (Abexa; 1:1000), rabbit anti-aromatase (in house antibody; 1:5000), rabbit anti-CVH (in house antibody 1:500), rabbit anti-FOXL2 (in house antibody; 1:2000), rabbit anti-p27 (Charles River Laboratories, 1:1000) and mouse anti-TOX3 (Novus Bio, 1:100). Sections were then washed in PBS and incubated for 1hr at room temperature with Alexa Fluor 488 donkey anti-Goat or Rabbit (1:1000) and Alexa Fluor 594 donkey anti-Rabbit or Mouse (1:1000) diluted in 1% BSA/PBS. Sections were washed, counterstained in DAPI/PBS and mounted in Fluorsave (Milipore). Images were collected on a Zeiss Axiocam MRC5.

### Tissue section fluorescence *in situ* hybridization

*CYP17A1* fluorescence *in situ* hybridization in E9.5 gonadal paraffin sections was performed as described before (23). To co-localize the steroidogenic cell marker *CYP17A1* with AMH, Aromatase or GFP, sections were subjected to antigen retrieval and then processed for immunofluorescence. We used goat anti-GFP 1:500, rabbit anti-aromatase 1:5000 or rabbit anti AMH 1:1000 and Alexa Fluor 488 donkey anti-Rabbit or Goat 1:1000 (secondary), followed by Sudan black to quench cell autofluorescence. Sections were counterstained in DAPI/PBS and mounted in Fluorsave (Milipore).

### RCAS plasmids generation

*TOX3* and *SOX9* open reading frames were amplified from chicken gonadal cDNA and Gibson cloned into the *RCASBP(D)-GFP-T2A* and *RCASBP(B)* viral vectors, respectively. *RCASBP(D)-GFP-T2A-TOX3* and *RCASBP(B)-SOX9* plasmid sequence were confirmed by sanger sequencing. Primers used can be found in Sup. table 1.

Two different shRNAs (*sh370* and *sh685*) were designed against *TOX3* ORF, ranked for effectiveness (75) and cloned into *RCASBP(A)-BFP plasmid* (37). Correct cloning and sequence were confirmed by Sanger sequencing. Primers used can be found in Sup. Table 1.

### DF-1 cell culture and transfection

DF-1 chicken fibroblastic cells were seeded onto 24 well plates and transfected with 1.5 µg of each construct according to the Lipofectamine 2000 protocol (Life Technologies). *RCASBP(A)-DMRT1* (19), *RCASBP(A)-SOX9* or *RCASBP(A)-GFP* plasmids were transfected in DF-1 cells in a 24 well plate. Forty-eight hours post transfection, cells were washed with 1X PBS, collected in 1ml of trizol and stored at -80°C till processing. To validate *TOX3* overexpression plasmid, *RCASBP(D)-GFP-T2A-TOX3* or *RCASBP(A)-GFP* were transfected in DF-1 cells. 48 hours post transfection cells were collected for RNA extraction or fixed for immunofluorescence against GFP and TOX3.

To test the shRNA, DF-1 cells plated in a 24 well plate were transfected with *RCASBP(A)-BFP-TOX3sh370, RCASBP(A)-BFP-TOX3sh685* or *RCASBP(A)-BFP-Firefly-Sh774* (non-silencing control) using Lipofectamine. After 48 hours, cells were transfected with *RCASBP(D)-GFP-T2A-TOX3*. 48 hours post transfection, cells were fixed briefly with 4% PFA in PBS and immunofluorescence against GFP was performed. To validate the ability of *TOX3 sh685* to knock down *TOX3* expression, T-25 flasks containing DF1 cells were transfected with *RCASBP(A)-BFP-TOX3sh685* or *RCASBP(A)-BFP-Firefly-Sh774* (non-silencing control). 72 hours post transfection they were collected and plated in 24 well plates. After resting for 24 hours, DF-1 cells were transfected with *RCASBP(D)-GFP-T2A-TOX3* overexpression plasmid. 48 hours post transfection, cells were collected in 1 ml of Trizol reagent and processed for RNA extraction.

### Virus purification

RCAS virus propagation and purification was performed as reported before (76). Briefly, viral plasmids were introduced into small T-25 flasks containing DF-1 cells using Lipofectamine 2000 (Life Technologies). Cells were passaged into six T-175 flasks and grown until they were super-confluent. Media was replaced with 1% FCS DMEM and harvested over 3 consecutive days. Virus was concentrated by ultracentrifugation, resuspended in 600 µl and titered.

### *In vivo* overexpression and knock down experiments

For *DMRT1* knockdown experiments *RCASBP(B)-GFP-DMRT1shRNA343* or *RCASBP(B)-GFP-scrambled* (control) live virus was injected into 0-day chicken blastoderms, as previously described (12). For *TOX3* knockdown experiments, *RCASBP(A)-BFP-TOX3sh685* or *RCASBP(A)-BFP-Firefly-Sh774* live virus was injected in 0-day chicken blastoderms. Embryos were collected at E9.5 in both cases.

*RCASBP(A)-DMRT1, RCASBP(A)-GFP or RCASBP(D)-GFP-T2A-TOX3* over-expression plasmids were electroporated *in ovo* into the left coelomic epithelium, as previously described (19). Embryos were collected at E7.5 for *DMRT1* overexpression experiments or E9.5 for *TOX3* overexpression experiments. The same methodology was used to introduce *RCASBP(A)-BFP-TOX3sh685* or *RCASBP(A)-BFP-Firefly-Sh774* knock down constructs. Embryos were also collected at E9.5.

