## Extended data for "DMRT1 regulation of *TOX3* modulates expansion of the gonadal steroidogenic cell lineage"

**Extended data Figure Legends:**

**Extended data 1. TOX3 sh685 reduced TOX3 expression in vitro**. (A) DF-1 cells expressing TOX3 sh370, sh685 or a non-silencing shRNA (NS shRNA) were challenged for 48 hours with RCASBP(D)GFP-T2A-TOX3 over-expression plasmid. The ability of the shRNA to knock down TOX3 was assessed by GFP intensity. (B) DF-1 cells expressing TOX3 sh685 or NS shRNA were challenged for 48 hours with RCASBP(D)GFP-T2A-TOX3 overexpression plasmid. TOX3 expression levels were quantified by qRT-PCR, relativized to β-actin and normalized to NS shRNA. Bars represent Mean ± SEM, n=6. *** = adjusted p value <0.001. Unpaired two-tailed t-test.

**Extended data 2. TOX3 knock down does not alter AMH and SOX9 expression**. TOX3 sh685 or NS shRNA (control) plasmids were electroporated in chicken E2.5 coelomic epithelium. Immunofluorescence detection of (A) DMRT1 or (B) AMH in E9.5 male gonadal sections. Dashed box indicates the magnified area. (C) Normalized expression of TOX3 and (D) CYP17A1 on a t-SNE visualization of all male gonadal chicken cells. Black arrows indicate the steroidogenic cell sub-cluster.

**Extended data 3. TOX3 over-expression in *vitro*.** (A) The TOX3 ORF was overexpressed in DF-1 cells. Immunofluorescence against GFP (transfection marker) and TOX3 was performed. White arrows indicate examples of TOX3 positive cells. (B) TOX3 or GFP (control) was overexpressed in DF-1 cells. TOX3 mRNA expression levels were measured by qRT-PCR. Expression is relative to β-actin and normalized to GFP overexpression control. Bars represent Mean ± SEM. * = p<0.05. Multiple t-test and Holm-Sidak post-test.

**Extended data 4. TOX3 over-expression in testis does not alter the fate of the supporting cell lineage.** TOX3 or GFP (control) over-expression plasmids were electroporated in chicken E2.5 coelomic epithelium. Male gonads were examined at E9.5 and immunofluorescence was performed to detect GFP and (A) AMH, (B) SOX9, (C) DMRT1 or (D) aromatase. Male gonads show normal expression of Sertoli cell markers AMH, SOX9 and DMRT1 and no aromatase expression.

**Extended data 5. TOX3 over-expression alters ovarian differentiation.** TOX3 or GFP (control) overexpression plasmids were electroporated in chicken E2.5 coelomic epithelium. Female gonads were collected at E9.5 and immunofluorescence was performed to detect GFP and (A) Cytokeratin, (B) CVH, (C) SOX9 or (D) DMRT1. Dashed box indicates the magnified area. Dotted line delineates the gonadal epithelium.

**Supplementary information:**

**Sup. Table 1.** List of primers used for cloning TOX3 and SOX9 overexpression and TOX3 shRNA (knockdown) RCASBP plasmids.

| Primer Name | Sequence |
|---|---|
| *TOX3 OE Fw* | *GAAAATCCCGGCCCCATGGATGTAAGATTTTACCCGTCCG* |
| *TOX3 OE Rv* | *GTCGTCCTTGTAGTCACTCAGAAAATACTGACCTGTGATAATACTTGAGTCTG* |
| *SOX9 OE Fw* | *GAGCTGACTCTGCTGGTGGCCTCGCGTACCACTGTGGCATATGAATCTCCTAGACCCCTTCATGAAAATGACAG* |
| *SOX9 OE Rv* | *GTCCCCTCAGATACGCGTATATCTGGCCCGTACATCGCATTTAAGGCCGGGTGAGCTGC* |
| *shRNA Fw* | *GGCCAGTGAATTCGCGTACCTCCTTCTCGCAGGGC* |
| *TOX3 sh370 Rv* | *AGCTATGACGAATTCGCAAAAAACCATTACTATATCCAGAAATCTCTTGAATTTCTGGATATAGTAATGGAAACCCCAGTTGCTCTCGG* |
| *TOX3 sh685 Rv* | *AGCTATGACGAATTCGCAAAAAAGATGCAGATGAATCAAATAGATCTCTTGAATCTATTTGATTCATCTGCATCAAACCCCAGTTGCTCTCGG* |

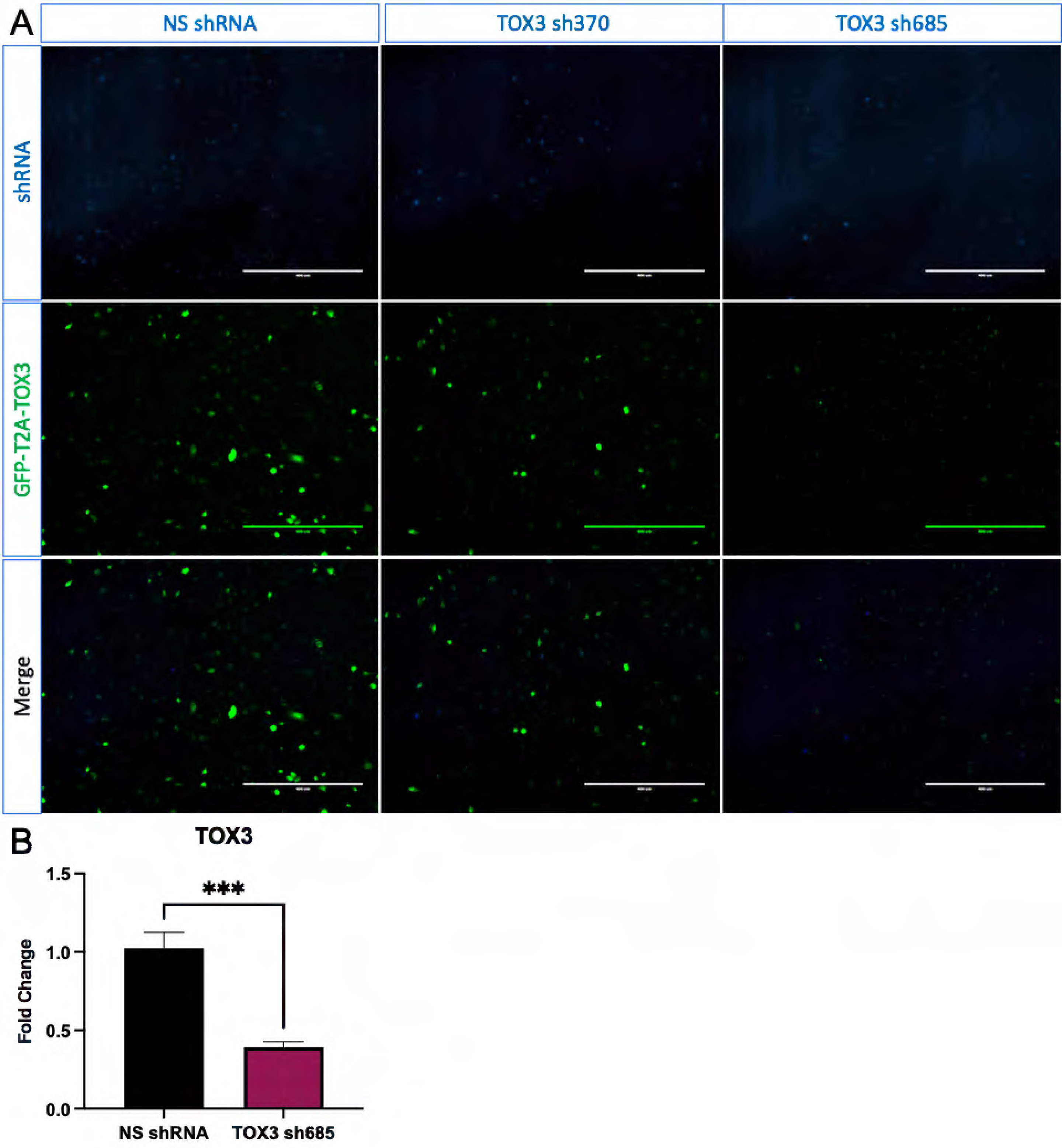
A
NS shRNA
TOX3 sh370
TOX3 sh685
shRNA
GFP-T2A-TOX3
Merge
B
TOX3
Fold Change
1.5
1.0
0.5
0.0
***
NS shRNA
TOX3 sh685

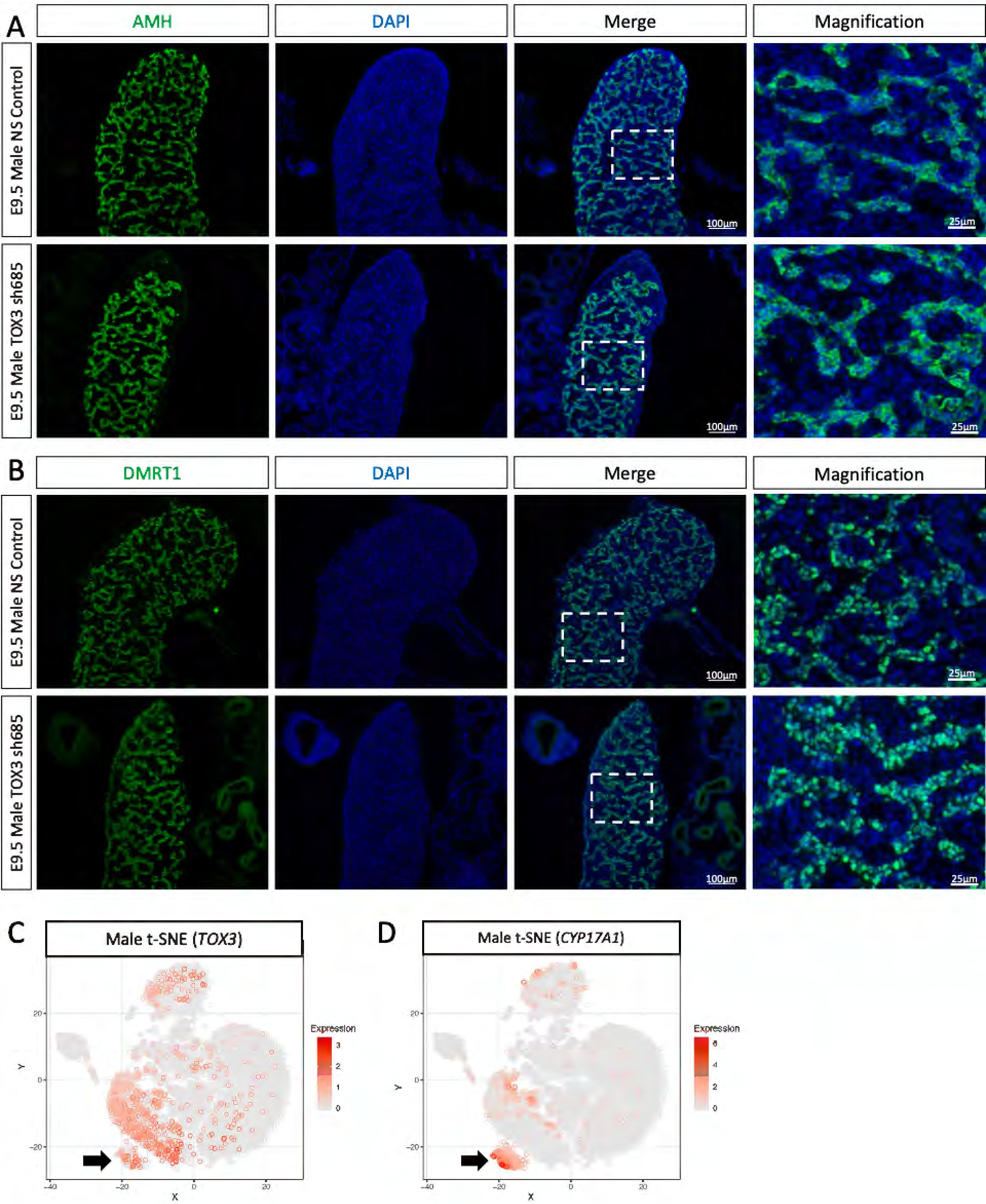
A
AMH
DAPI
Merge
Magnification
E9.5 Male NS Control
E9.5 Male TOX3 sh685
100µm
25µm
B
DMRT1
DAPI
Merge
Magnification
E9.5 Male NS Control
E9.5 Male TOX3 sh685
100µm
25µm
C
Male t-SNE (TOX3)
Expression
D
Male t-SNE (CYP17A1)
Expression
X
Y

A

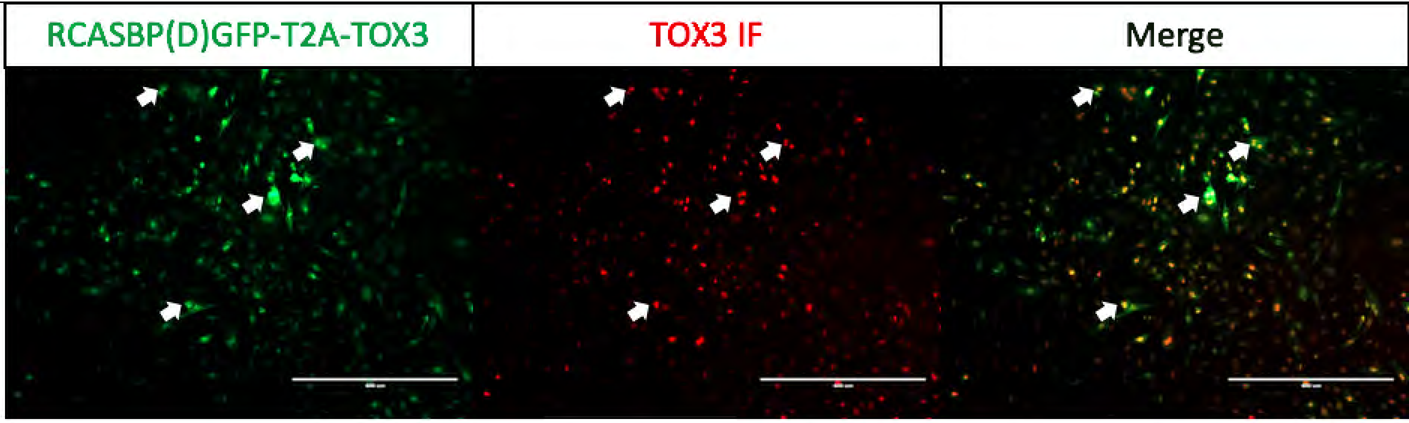
RCASBP(D)GFP-T2A-TOX3
TOX3 IF
Merge

B

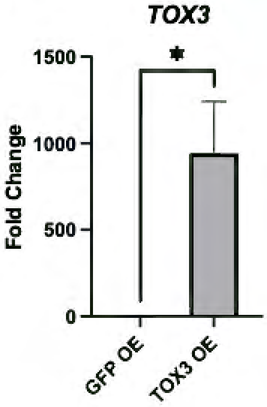
TOX3
*
Fold Change
1500
1000
500
0
GFP OE
TOX3 OE

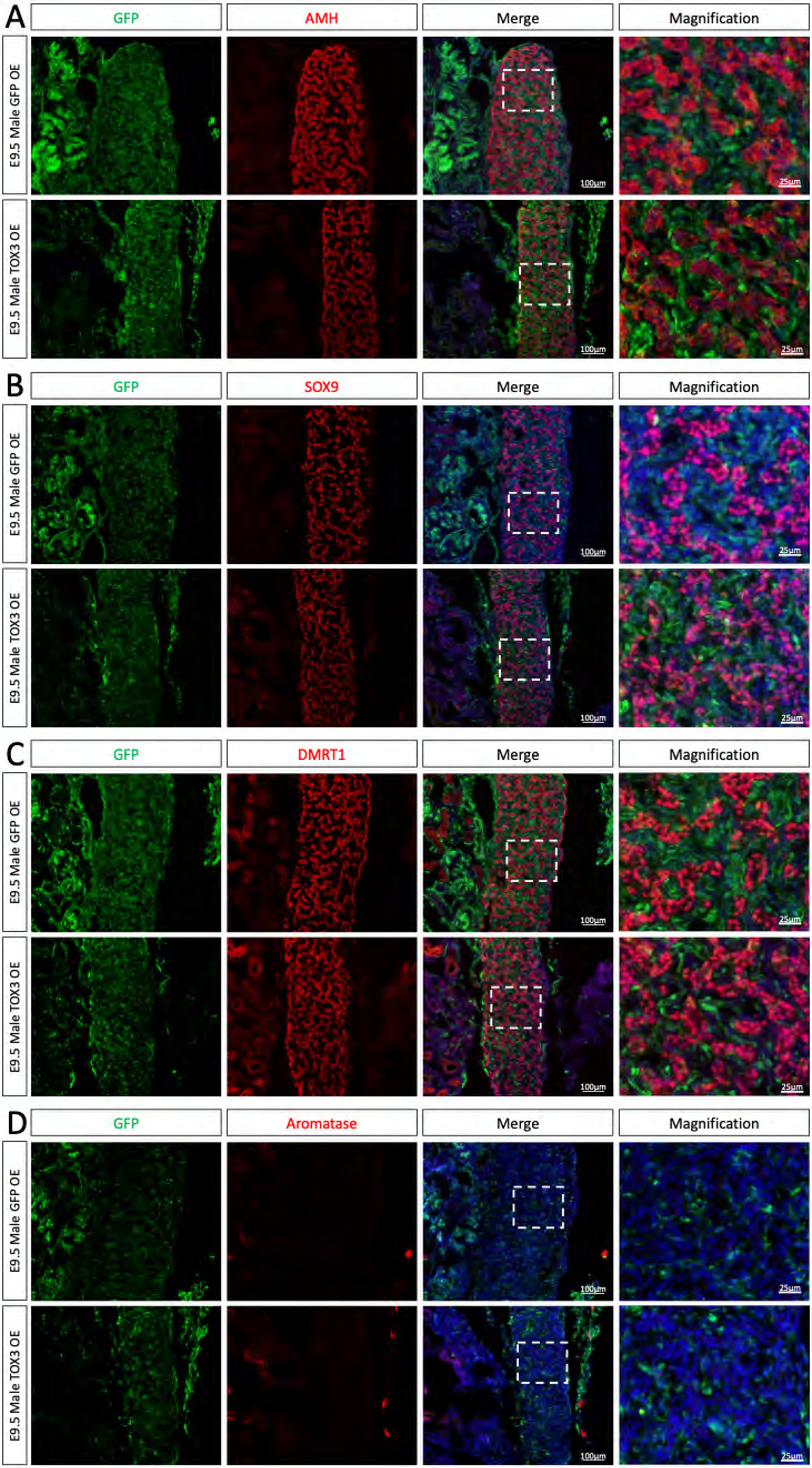
A
GFP
AMH
Merge
Magnification
E9.5 Male GFP OE
E9.5 Male TOX3 OE
100μm
25μm
B
GFP
SOX9
Merge
Magnification
E9.5 Male GFP OE
E9.5 Male TOX3 OE
C
GFP
DMRT1
Merge
Magnification
E9.5 Male GFP OE
E9.5 Male TOX3 OE
D
GFP
Aromatase
Merge
Magnification
E9.5 Male GFP OE
E9.5 Male TOX3 OE

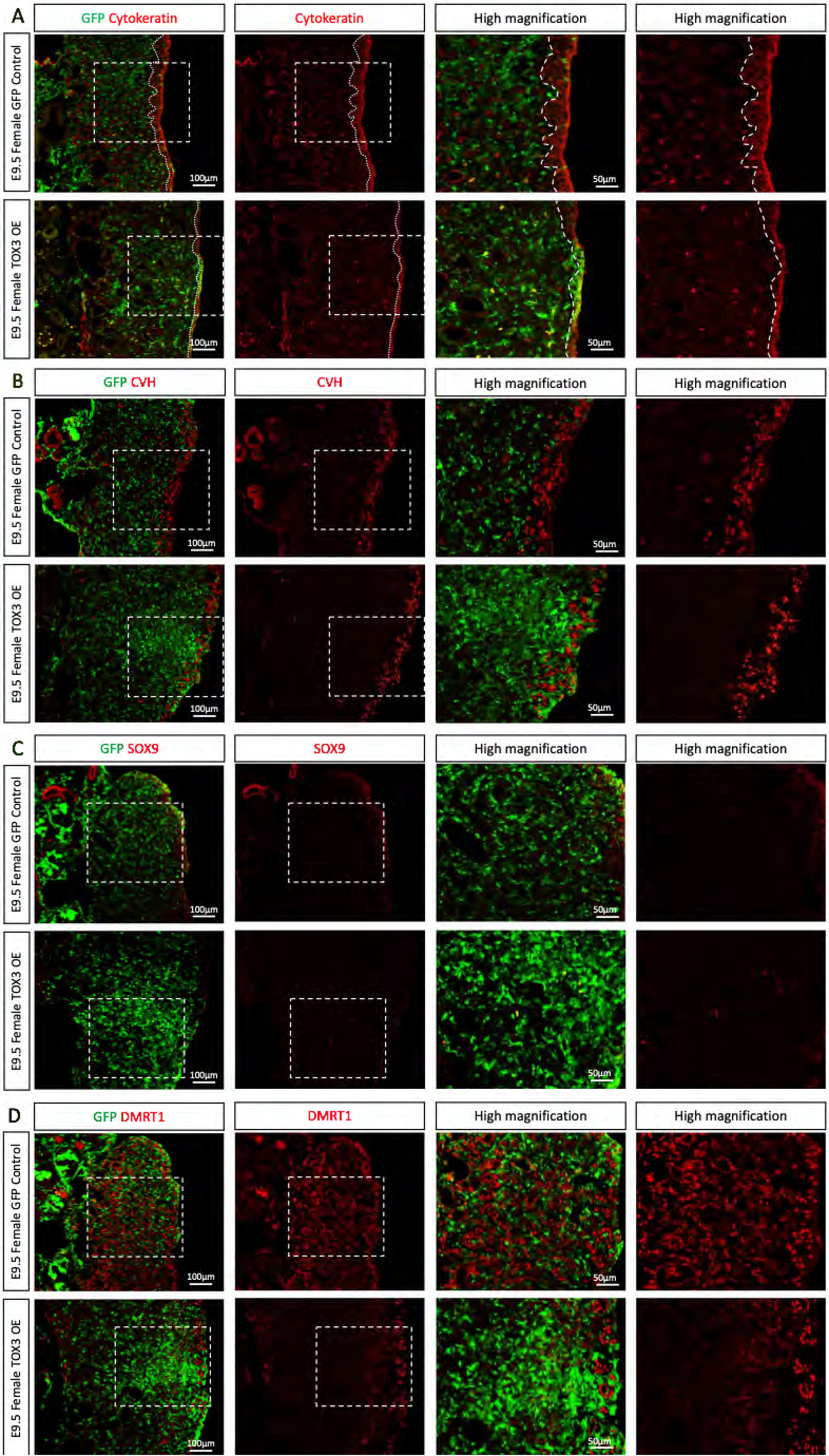

A
GFP Cytokeratin
Cytokeratin
High magnification
High magnification
E9.5 Female GFP Control
E9.5 Female TOX3 OE
100μm
50μm
B
GFP CVH
CVH
High magnification
High magnification
E9.5 Female GFP Control
E9.5 Female TOX3 OE
100μm
50μm
C
GFP SOX9
SOX9
High magnification
High magnification
E9.5 Female GFP Control
E9.5 Female TOX3 OE
100μm
50μm
D
GFP DMRT1
DMRT1
High magnification
High magnification
E9.5 Female GFP Control
E9.5 Female TOX3 OE
100μm
50μm
